# Evolutionary processes make invasion speed difficult to predict

**DOI:** 10.1101/013680

**Authors:** Ben L. Phillips

## Abstract

A capacity to predict the spread rate of populations is critical for understanding the impacts of climate change and invasive species. Despite sophisticated theory describing how populations spread, the prediction of spread rate remains a formidable challenge. As well as the inherent stochasticity in the spread process, spreading populations are subject to strong evolutionary forces (operating on dispersal and reproductive rates) that can cause accelerating spread. Despite these strong evolutionary forces, serial founder events and drift on the expanding range edge mean that evolutionary trajectories in the invasion vanguard may be highly stochastic. Here I develop a model of spatial spread in continuous space that incorporates evolution of continuous traits under a quantitative genetic model of inheritance. I use this model to investigate the potential role of evolution on the variation in spread rate between replicate model realisations. Models incorporating evolution exhibited more than four times the variance in spread rate across replicate invasions compared with nonevolving scenarios. Results suggest that the majority of this increase in variation is driven by evolutionary stochasticity on the invasion front rather than initial founder events: in many cases evolutionary stochasticity on the invasion front contributed more than 90% of the variance in spread rate over 30 generations. Our uncertainty around predicted spread rates - whether for invasive species or those shifting under climate change - may be much larger than we expect when the spreading population contains heritable variation in rates of dispersal and reproduction.

## 2 Introduction

Predicting the spread rate of biological invasions has been a longstanding interest of ecologists (Skellam, 1951; Elton, 1958; Hengeveld, 1989). Predicting spread rate is useful for the management of invasive species, but is also fundamental to understanding the dynamics of range shift in response to climate change, both past and present (Shigesada and Kawasaki, 1997; Clark et al., 1998; Sax et al., 2005). Increasingly, it is also appreciated that understanding the dynamics of spread may also be important to medicine; in understanding tumour growth and the formation of biofilms (Orlando et al., 2013; van Ditmarsch et al., 2013). Despite these new applications, the long-standing interest of ecologists, and the development of elegant and sophisticated theory around the dynamics of spreading populations (e.g., Hastings, 1996), our capacity to forecast spread rate remains poor (Hastings et al., 2005).

There is, perhaps, little surprise in this observation. The process of range shift results, after all, from the interplay of two stochastic processes intrinsic to the population: dispersal and population growth. These intrinsic processes are, in turn, affected by spatial and temporal variation in the environment. Thus, we should not be surprised that there will always be fundamental limits to our capacity to forecast spread rate. Indeed, even in controlled environments, replicated invasions exhibit wildly varying spread rates (Melbourne and Hastings, 2009).

In recent years it has also become apparent that some of our failure to predict spread rate - particularly in situations of accelerating spread – is likely the result of rapid evolution (Travis and Dytham, 2002; Phillips et al., 2008). The process of spread creates powerful selective forces that favour individuals on the invasion front with higher dispersal and reproductive rates (Phillips et al., 2010; Shine et al.. 2011; Benichou et al., 2012; Kubisch et al., 2013). These selective forces can, on ecologically-relevant timescales, alter phenotypes on invasion fronts resulting in accelerating spread (Perkins, 2012; Perkins et al., 2013).

As well as the realisation that rapid evolution can cause profound shifts in spread rate, we are also beginning to appreciate that evolution on invasion fronts can also be highly stochastic (Excoffier and Ray, 2008; Excoffier et al., 2009). The process of range shift creates a situation of serial founder events — only relatively few individuals make up the invasion vanguard each generation — that can see even maladaptive alleles ‘surf’ to high frequency and be smeared across the invaded range (Klopfstein et al., 2006; Travis et al., 2007). This situation of serial foundering is akin to genetic drift: it is drift through space rather than time (Slatkin and Excoffier, 2012). The resulting accumulation of deleterious alleles on invasion fronts has recently been termed ’’expansion load” and is currently the focus of intense theoretical interest (Peischl et al., 2013).

Thus, rapid evolution can drive important variation in spread rate, but the ultimate evolutionary trajectory of a spreading population is subject to a high degree of stochasticity through both an initial founder event at colonisation and then serial founder events happening on the invasion front every generation. It remains possible, therefore, that this evolutionary stochasticity contributes to the variation in spread rates we see in nature and in replicated invasion in the lab. To test this idea, I develop an individual-based simulation model that allows both dispersal and reproductive rate to evolve. Analysis of this model suggests that evolutionary process can lead to a massive increase in the variability of realised spread rates.

## 3 Methods

To examine the contributions of various stochastic forces to variation in range shift, I develop a discrete-time simulation model of population and evolutionary dynamics. The model tracks sexually hermaphroditic individuals that shift and reproduce over time in continuous space. I use a quantitative genetics model for the inheritance of two continuous traits: one affecting dispersal and the other affecting fitness. The model is developed and analysed primarily in 1-dimensional space, with a brief extension to the 2-dimensional case. All numerical procedures were conducted in R (R Development Core Team, 2012), and the code is available as an online supplement to this paper (Appendix S2).

Individuals express two phenotypes, one relevant to dispersal *z*_*d*_, and one relevant to fitness *z*_*w*_. Both traits are continuous (both ∈ ℝ) with an underlying quantitative genetic structure that assumes trait genetic covariances are zero (there is no genetic correlation between the traits). By taking the individual-based approach, demographic stochasticity and the evolution of trait means and variances can be easily incorporated. Because it is impractical to completely explore parameter space in such a model, however, I take the approach of providing a proof-of-concept; demonstrating the potential importance of evolutionary process in generating both rapid and less predictable invasion speeds. An analytical treatment of these ideas remains a formidable challenge.

### 3.1 Population dynamics

Generations are discrete and non-overlapping. Local dynamics are determined by reference to a continuous spatial “neighbourhood” whose size remains constant. Both the density of individuals and the mean and variance of trait values in an individual’s neighbourhood are calculated as a sum or mean across the entire population weighted for distance from the individual. The weighting through space is simply a Gaussian probability density function with a standard deviation set to one. Thus, local density at location *x*, *n*_*x*_, is given as a sum over the *p* individuals in the population:

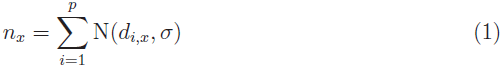

Where N() is the Gaussian pdf, *d*_*i*,*x*_ is the distance between individual *i* and location *x*, and *σ* is the smoothing scale (a constant; standard deviation = 1 in this case). The smoothing scale (the scale of local dynamics) is independent of dispersal values in the population. This choice effectively sets up a reproductive phase (in which local interactions matter) and a dispersal phase (in which local interactions do not matter). The smoothing scale, in conjunction with the population carrying capacity (*n*\*, see below) together define the approximate neighbourhood size (= 4*n*\**σ*) of demes within the population.

Individual reproduction is a stochastic density-dependent process. I calculate an expected number of offspring for each individual E(*o*_*i*_) using the Beverton-Holt (Beverton and Holt, 1957) population growth function 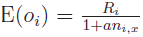 Where *R*_*i*_ is an individual’s expected density-independent fecundity (determined by its *z*_*w*_ phenotype, see below), *n*_*i*,*x*_ is the density at the individual’s location, and *a* is a constant that determines the strength of intraspecific competition. At the beginning of simulations, *a* is set to 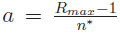 where *R*_*max*_ is the maximum expected fecundity achievable by an individual in the population, and *n*^*\**^ is the carrying capacity that would be achieved if all individuals achieved a fecundity of *R*_*max*_.

A subset of simulations also involved a strong Allee effect. To achieve the Allee effect, I simply multiplied *R_i_* by 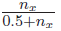, which produces inverse density dependence at low population densities (Stephens and Sutherland, 1999). Irrespective, once an expected number of offspring is calculated, the realised number of offspring is drawn from a Poisson distribution with mean of E(*o*_*i*_). Beverton-Holt dynamics have the useful property that population dynamics remain stable irrespective of the population growth rate. This choice of local dynamic ensures that local population fluctuations are purely due to demographic stochasticity.

Following reproduction, all parents die and offspring disperse. Individuals disperse according to a draw from a Gaussian distribution with a mean of zero and a standard deviation given by *e*^*z*_*d*,*i*_^, where *z*_*d*,*i*_ is the individual’s dispersal phenotype.

### 3.2 Trait evolution

Individuals express two phenotypes, one, *z*_*w*_, that determines the individual’s maximum expected reproductive rate, and the other, *z*_*d*_, that determines the individual’s dispersal propensity. An individual’s reproductive rate is influenced by how well adapted it is to the local environment, where the optimum value for *z*_*w*_ is 0. The density-independent fecundity of an individual, *R*_*i*_, is a function of an individual’s *z*_*w*_ as,

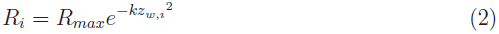

where *R*_*max*_ is the upper limit on expected individual fecundity (constant across individuals), and *k* is a constant defining the strength of stabilising selection in the system (here set to a constant value of 2).

The inheritance of traits is determined by a basic quantitative genetic model in which individuals mate with a partner drawn at random (with probability weighted by *N*(*d*_*i*,*x*_, *σ*)) from the population. All individuals reproduce and each individual carries a “breeding value” that contributes to the phenotype of its offspring. In quantitative genetics, an individual’s breeding value represents the sum of the additive genetic contributions to its phenotype and is usually estimated from the mean of its offspring (Lynch and Walsh, 1998). Instead of estimating breeding values, I define them here for each individual before its offspring are produced. Importantly, the variance in breeding values in a randomly mating population is identical to the additive genetic variance in that population (Lynch and Walsh, 1998), so initially, each individual’s breeding value can be determined by a draw from a normal distribution with mean set to the trait mean and variance equal to the additive genetic variance of the population *V*_*a*_ (see below). Offspring breeding values are then centered on the resulting mid-parent breeding value (the mean of the breeding values of the parents), but deviate from this mid-parent value according to a normal distribution with variance equal to half the variance in local mean breeding values (i.e., half the local additive variance). This distribution of offspring breeding values is that expected in a situation with no dominance or epistasis (Roughgarden, 1979), and largely reflects variance between offspring driven by segregation (i.e., randomness in the set of parental alleles inherited by each offspring). Thus, for example, the breeding value for dispersal of offspring one from parent one at location *x* is calculated as,

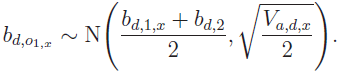

Local trait mean breeding values (*b*̅_*d*,*x*_ and *b*̅_*w*,*x*_) and variances are calculated over the same spatial neighbourhood as that governing local dynamics. For example (for bd),

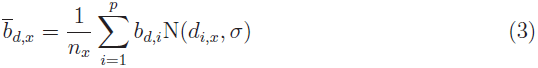

and local trait genetic variances are calculated as,

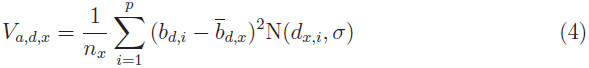

Thus, I only track standing variation: genetic variances evolve through space as a consequence of gene flow (movement of individuals), and are eroded by selection, but are not subject to the additional inflationary force of mutation. The modelled scenario focusses on small population sizes and short-term dynamics in which mutational contributions to variance are likely to be negligible. The offspring’s final phenotype (*z*_*d*_, *z*_*w*_) is determined by adding environmental variance (*V*_*e*_) to the breeding values again using a random draw from a Gaussian distribution. For example,

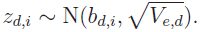

*V*_*e*_ is constant over time and space: when the model is initiated, the amount of environmental variance added to offspring breeding value is determined according to the trait’s heritability *h*^2^ and total phenotypic variance, *V*_*p*_, where *V*_*p*_ = *V*_*e*_ + *V*_*a*_ and *V*_*a*_ = *h*^2^*V*_*p*_ and.

To keep spread rate modest (and computationally tractable), I set the initial phenotypic mean of *z*_*d*_ to ln(4) (a root mean square displacement of 4 units) and the initial total phenotypic variance of *z*_*d*_ to 0.2 in all simulations. At initialisation, then, the average individual will leave it’s local neighbourhood (defined as ±2 units from its location). The initial mean *z*_*w*_ phenotype was set to zero (the optimum value) and again total phenotypic variance was set to 0.2.

### 3.3 The modelled scenario

All simulations begin with the introduction of 20 individuals to a point in space, and the model examines variation in realised spread distances over 30 generations. For the 1-dimensional case, I examined values of *n*\* between 10-50, and *R*_*max*_ between 2-20. At the end of these 30 generations, the distance spread (the distance between introduction and the location of the furthest individual) is recorded for both invasion fronts (Fig. 1). Recording spread distance of the twin invasion fronts allows me to calculate a within-population measure of variation in spread rate.

**Figure 1.**
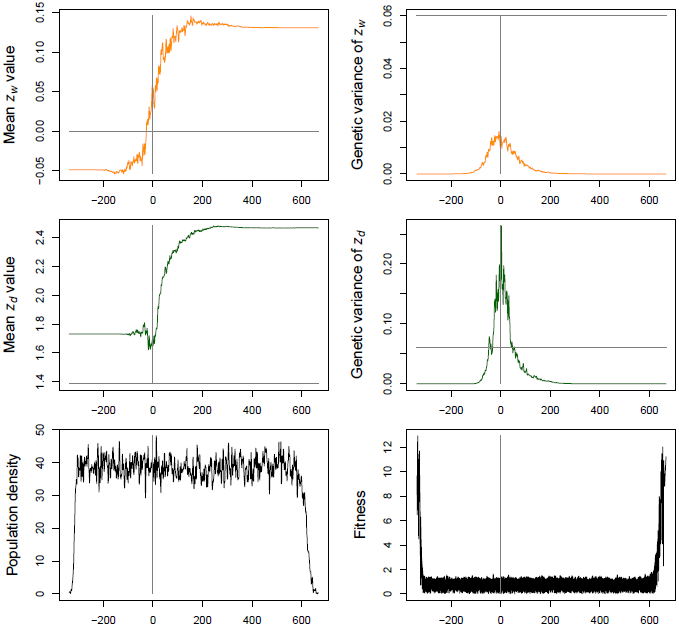
A typical realisation of the model in 1-dimension following 30 generations of spread. Grey lines represent initial trait value (horizontal line) or introduction location (vertical line). First row: mean and genetic variance of the trait affecting reproductive output (*z*_*w*_). Initial trait value in this case is zero, and the figure indicates substantial drift from this optimum, moderate erosion of genetic variance at the introduction locale, and substantial loss of genetic variance towards the invasion front. Second row: mean and genetic variance of the trait affecting dispersal (*z*_*d*_). In this case the two invasion fronts have evolved very different values for the trait, trait variance has increased from initial values at the introduction location, but still decreases strongly towards the invasion front. Third row: population density through space, and mean individual fitness E̅(*o*) through space. The genetic neighbourhood of each individual spans an interval of four units along the x-axis.

For each set of parameters examined, I created 20 founder populations (each containing 20 individuals). For each founder population of 20 individuals, 20 replicate realisations of an invasion were undertaken: ten with trait heritabilities set to zero, and ten with trait heritabilities set to 0.3. For each of these replicates, spread distance was recorded for both invasion fronts. Initial trait variances were kept identical between the evolving and non-evolving scenarios (although trait variances then evolved in the evolving scenario as *V*_*a*_ evolves). With this arrangement of replicated invasions, I can partition variation in spread distance into that resulting from,

1. stochasticity in dispersal and demography (the total variance – within and across populations – in spread distance in the non-evolving populations), *S*_0_. Noting that founder events do not occur in the non-evolving scenario, because the traits are not heritable;
2. initial founder events (the between-population variance in spread distance in the evolving populations), *S*_*f*_; and
3. stochasticity in the evolutionary process (the within-population variance in spread distance in the evolving populations, *S*_*e*_, minus that due to pure demographic stochasticity, *S*_0_.

The 2-dimensional case was substantially more computationally intensive, so for this case I only examined one value of *n*\* (*n*\* = 10), and only examined a smaller range of *R*_*max*_ (between 2-10). I otherwise kept the design the same as the 1dimensional case (Fig. 2). The other difference for the 2-dimensional simulations was that measurements of spread distance were taken along both the *x*, and *y* axes: individuals’ locations in (*x*, *y*) were collapsed onto each axis yielding two sets of twin invasion fronts. Other than generating twice as many within-population measures of spread rate, subsequent analysis proceeded as per the 1-dimensional case.

**Figure 2.**
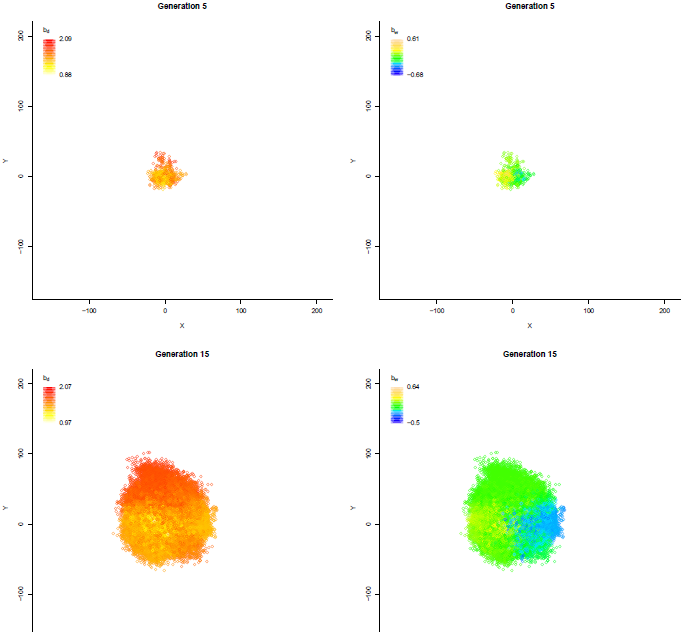
A typical realisation of the model in 2-dimensions. Snapshots at time 5, 15, and 30 generations are shown at each row of the figure panel. The columns of the figure panel correspond to the traits for dispersal *z*_*d*_ and fitness *z*_*w*_ respectively. Colours allow us to observe individual breeding values for these two traits and how these vary over space and time. Note that, like the 1-dimensional case, different evolutionary outcomes emerge on different parts of the invasion front. In the present case higher dispersal values emerge in the upper sector, and lower fitness values emerge on both right and left sectors. Together, these differing evolutionary trajectories lead to differing spread rates (clearly apparent in the lower panel, in which a circle, and crosshairs centred on the starting position, have been placed for reference).

## 4 Results

### 4.1 1-dimensional situation

The model deliberately investigated very short time periods (30 generations of spread) and focussed on realistic trait heritabilites (usually = 0.3). Despite the short spread time and the modest heritability (=0.3), models with heritable traits resulted in greater spread rates (Fig. 3). When heritability was 0.3, spread rates were, on average across all simulations, 1.3 times faster. Although evolution resulted in spread rates 1.3 times faster than the non-evolving scenario, the variance in spread rate was, on average, 4.2 times larger in the evolutionary scenarios (Fig. 3). Importantly, much of this additional variation manifested as variation within a single realisation of spread: i.e., large differences in the spread rate between the twin invasion fronts (see e.g., Fig. 1). That is, there were typically differences in the evolutionary trajectory on the twin invasion fronts.

**Figure 3.**
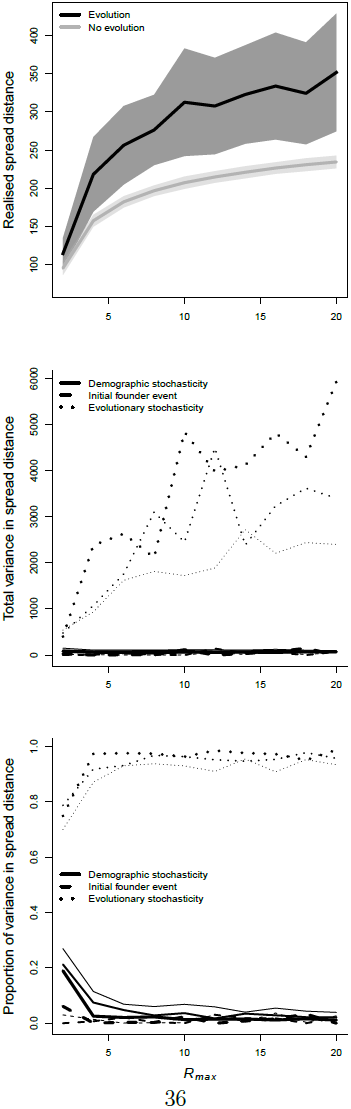
Spread distances in a 1-dimensional space under scenarios with and without trait evolution. The top panel shows spread distance with and without trait evolution across a range of *R*_*max*_ (with *n*\*=50). Shaded areas represent the standard deviation in spread distance across replicate invasion fronts. It is clear that allowing dispersal and reproductive rate to evolve both increases the rate of invasion as well as the variation in that rate across realisations. The second panel shows variance in spread distance decomposed into that resulting from demographic stochasticity, initial founder events, and evolutionary stochasticity. Line thickness in lower two panels represents different values of infraspecific competition (thick to thin; *n*\* = 50, 25, 10 respectively). The lower panel shows this same decomposition of variance as a proportion of total variance. At low rates of population increase (*R*_*max*_ = 2) demographic stochasticity contributes around 20% of the variance in spread distances in the model. Above these low values of *R*_*max*_, however, stochastic evolutionary processes on the expanding range edge (serial foundering: ‘Evolutionary stochasticity’) accounts for most of the variation in spread distance.

Why did these differences emerge? In this model, the signal of phenotype surfing (driven by serial founder events) was evident in the fitness trait, which often showed substantial deviations from optimal values (i.e., zero) on the advancing range margins. That is, serial founder events clearly undermine the stabilising selection operating on this trait. This same phenotype surfing must also have occurred for the dispersal trait, although this trait was subject to strong directional evolution (driven by spatial sorting and natural selection, see Phillips et al. (2010)), and so in this case surfing manifests as differing rates of evolutionary shift. Acting on both traits together, serial founder events (“surfing”) resulted in strong disparities in the evolved spread rate on each of the twin invasion fronts. The “final” dispersal and fitness values on each front will determine the equilibrium spread rate (Fisher, 1937; Benichou et al., 2012), so these varying evolutionary trajectories caused substantial variation in spread rate between twin invasion fronts.

Overall, this evolutionary stochasticity typically contributed to most (often more than 90%) of the variance in spread rate in the 1-dimensional model (Fig. 3), with much lower levels of variance being attributable to pure demographic stochasticity and the initial founder effect (Fig. 3). This basic result appears reasonably robust to variation in the equilibrium population density (*n*\*) and maximum fecundity (*R*_max_, Fig. 3). Demographic stochasticity did, however, play a larger role when intrinsic rates of population growth (*R*_*max*_) were low, and when trait heritability was low (Appendix S1). Interestingly, the inclusion of a strong Allee effect lowered the overall stochasticity in spread rate, but did not greatly alter the proportional contributions of demography, initial founder effect, and evolution (Appendix S1).

### 4.2 2-dimensional situation

Here again, models with heritable variation resulted in faster spread rates (Fig. 4). This increased spread rate for the 2D case was similar to that for the 1D case. If we compare the overall increase in spread due to evolution between the two spaces (bearing in mind that we can only use a subset of the 1D cases for comparison: *R*_*max*_ = {2, 4, 6, 8,10} and *n*\* = 10) then the ratio of proportional increase in spread rate 1D:2D is 1.21:1.16. As with the 1D case, overall variance in spread rate was substantially higher for the evolved scenario (Fig. 4) but, this increase was more modest in the 2D case compared with the equivalent parameter settings in the 1D case: the ratio of proportional increase in variance, 1D:2D was 2.52:1.84.

**Figure 4.**
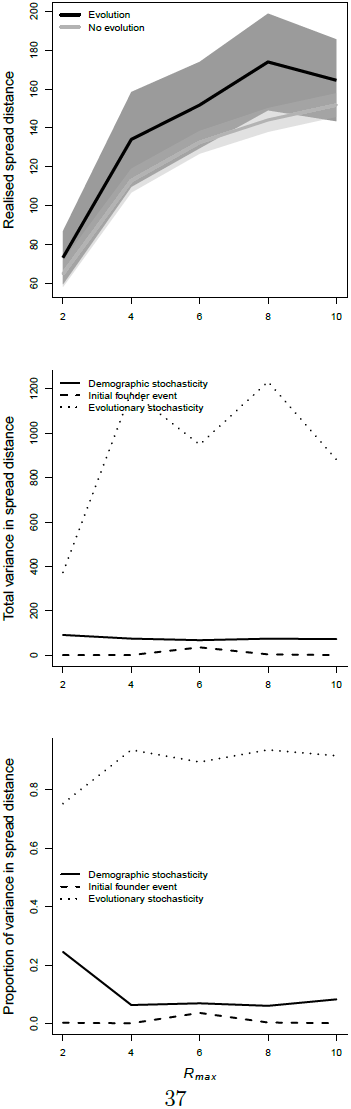
Spread distances in a 2-dimensional space under scenarios with and without trait evolution. The top panel shows spread distance with and without trait evolution across a range of *R*_*max*_ (with *n*\* =10). Shaded areas represent the standard deviation in spread distance across replicate invasion fronts. It is clear that allowing dispersal and reproductive rate to evolve both increases the rate of invasion as well as the variation in that rate across realisations. The second panel shows variance in spread distance decomposed into that resulting from demographic stochasticity, initial founder events, and evolutionary stochasticity (again *n*\* =10 throughout). The lower panel shows this same decomposition of variance as a proportion of total variance. At low rates of population increase (*R*_*max*_ = 2) demographic stochasticity contributes around 20% of the variance in spread distances in the model. Above these low values of *R*_*max*_, however, stochastic evolutionary processes on the expanding range edge (serial foundering: ‘Evolutionary stochasticity’) accounts for most of the variation in spread distance.

As with the 1D case, much of this increased variance in spread rate manifested as variation within a single realisation of spread. That is, there were substantial differences in spread rate between different sectors of the invasion and these differences were driven by differing evolutionary trajectories on various parts of the invasion front (Figs 2 and 4). Thus, although, the overall increase in variation in spread rate was smaller in the 2D case compared with the 1D case, the proportional contributions of demography, initial founder event, and evolutionary stochasticity, were similar: evolutionary stochasticity typically generated more than 90% of the observed variation in spread rate.

## 5 Discussion

If we are to accurately forecast rates of spread, whether for toads, tumours, or trees, it is clear that we need to account for variation in dispersal and reproduction (e.g., Neubert et al., 2000; Schreiber and Ryan, 2011). In purely ecological models of spread (i.e., those that do not incorporate evolution) demographic stochasticity can act to either slow (e.g., Lewis, 2000; Snyder, 2003) or increase (e.g., Ellner and Schreiber, 2012) spread rates relative to deterministic expectations. Evolutionary models of spread, on the other hand, typically lead to higher rates of spread than those predicted by equivalent ecological models because dispersal and reproductive rates evolve upwards during range advance (Burton et al., 2010; Phillips et al., 2010; Perkins et al., 2013). My analysis now suggests that evolutionary processes not only make invasions faster, they also make invasion speed more unpredictable, because very different spread rates can emerge from identical starting conditions.

It appears that much of this variation in spread rate might be due to evolutionary stochasticity: evolutionary processes pushing through a strong stochastic filter. This evolutionary stochasticity is a result of the serial founder events that occur on the invasion front each generation. Clearly, the analysis here is not exhaustive: an infinite parameter space exists within the model and exploring it all is not practicable. Thus, I aim mostly for a proof-of-concept: to demonstrate that evolutionary stochasticity might play an important role in making invasions inherently difficult to forecast, and that it can do so across a wide range of circumstances. The work complements recent work by Peischl et al. (2015) which shows that the accumulation of fitness-reducing variants on an expanding range edge can limit the rate of spread. Here, I show that both dispersal and fitness traits are affected by serial foundering, and the consequent evolutionary stochasticity in spread rate is substantial. Indeed, the large role of evolutionary stochasticity appears to be a robust result: evolutionary stochasticity is associated with more than 90% of the variation in spread rate, in both one and two dimensions and across a large range of reproductive rates, equilibrium densities, and heritabilities. By contrast, other processes acting alone — demographic stochasticity and initial foundering — often account for relatively minor proportions of the overall variance in spread rate (Figs 3 and 4).

Clearly, however, the degree to which evolutionary stochasticity makes invasions unpredictable will depend on the system at hand. One constraint with this system is that, for logistical reasons, the population densities I have used are relatively modest (at *n*\* = 50, for example neighbourhood size is approximately 200 individuals with continuous gene flow between demes), and so we might expect drift to be an important force in this system, even at equilibrium. Nonetheless, varying *n*\* made little difference to the proportion of variance attributable to evolutionary stochasticity (see lower panel of Fig. 3). This is exactly what we would expect if evolutionary dynamics were dominated by conditions on the leading edge of the invasion. Population densities on this leading edge are low and serially foundered irrespective of the equilibrium density; so varying *n*^*^ makes little difference to the degree of drift on the invasion front.

The model here also shows that absolute variance in spread rate is much larger in a 1D space compared with the 2D case. Stochastic effects are exaggerated in the 1D case because serial founder events are not moderated by the spatial autocorrelation that acts across another spatial dimension. In other words the effect of drift on the invasion front is lower in the 2D case. As well as reducing the strength of the serial founder effect, in two dimensions, gene flow perpendicular to the direction of invasion should eventually ensure that a close-to-optimal invasion phenotype emerges across the entire invasion front (see Hallatschek et al., 2007). But although this should happen, it will take time. In two dimensions, and given sufficient time, we would expect different rates of invasion to emerge across different sectors of the invasion.

These different rates will remain stable for a period of time before being invaded (from the side) by sets of phenotypes that generate more rapid invasion speed (a phenomenon apparent in Fig 2).

Another spatial dimension might also alter the evolutionary effect by slowing down the erosion of genetic variance that occurs very rapidly in the 1D case. A slower loss of variance would likely allow the stochastic aspects of the evolutionary process more time to play out; giving vanguard populations more time to drift. The decay of genetic variance could also be slowed by allowing greater capacity for longdistance dispersal. Long-distance dispersal, because it creates greater mixing, has been shown to slow the loss of genetic variance on invasion fronts (e.g., Bialozyt et al., 2006). Thus, swapping the Gaussian dispersal kernel used here for one with fatter tails would slow the loss of genetic variance on the invasion front and, because of this, potentially increase the proportion of variance in spread rate contributed by evolutionary stochasticity. More generally, the shape of the dispersal kernel is already known to have a major impact on invasion dynamics (Kot et al., 1996). Highly leptokurtic kernels, for example, can result in patchy and highly stochastic invasions (Lewis and Pacala, 2000; Bocedi et al., 2014) — a situation where serial founder events might be particularly pronounced. Thus, the shape of the dispersal kernel remains a critical consideration for both the ecological and evolutionary dynamics of invasion.

The model I explore also treats the environment as homogenous in space and time. Adding stochastic environmental variation to the model would presumably increase the role of “demographic” stochasticity in generating variance in spread rates. Even so, it is likely that evolutionary stochasticity will still play a strong role in making spread rate unpredictable. The reason for this is that evolution generates autocorrelated variation in dispersal and demographic rates: this autocorrelation emerges in the evolutionary model simply because traits are heritable. The work of Schreiber and Ryan (2011) clearly shows that autocorrelated demographic stochasticity increases the overall unpredictability of spread rate above that seen when stochasticity is not autocorrelated. Thus, even in a model with strong environmental stochasticity, we would expect evolutionary effects to make spread rate even more unpredictable.

Finally, the introduction of an Allee effect reduces both evolutionary and demographic stochasticity. By forcing the vanguard population to grow from a larger founder population each generation, Allee effects reduce demographic stochasticity and also slow the loss of genetic variation in the vanguard (Taylor and Hastings, 2005; Hallatschek and Nelson, 2008). As well as this, by creating a negative correlation between fitness and dispersal on the invasion front, Allee effects can undermine the evolutionary processes that lead to increasing dispersal and reproductive rates (e.g., Travis and Dytham, 2002; Phillips, 2009; Burton et al., 2010). As a consequence, Allee effects may play a particularly powerful role in decreasing overall stochasticity during invasion. Indeed, when I incorporated an Allee effect, the model showed much lower overall stochasticity in spread distance although, again, the proportion of this variance due to evolutionary stochasticity remained high (Appendix S1).

There are, of course, other ways in which the role of evolutionary stochasticity during invasion can be diminished. Most obviously, if there is no genetic variance for the traits that determine spread rate (reproductive and dispersal rates), evolutionary stochasticity will not be an issue. Where there is even a small amount of heritable variance in dispersal and reproductive rates, however, my analysis suggests that evolutionary stochasticity can play a powerful role in generating unpredictable invasion speeds. Obviously, extrinsic factors (e.g., environmental heterogeneity, or species interactions) will still play an important role (e.g., Tobin et al., 2007; Burton et al., 2010), but the analysis here suggests that a great deal of variance in spread rate may be explainable by intrinsic properties of the population, without the need to invoke variation in the environment. This observation is at once encouraging — environmental variation may be less important than we currently think — and discouraging, because to be clear about our predictive uncertainty requires information on population attributes that are difficult to measure (e.g., trait heritabilities). What is clear, however, is that evolutionary processes — both deterministic and stochastic — are of critical importance in predicting invasion speed and in understanding our uncertainties around these predictions.

## 6 Acknowledgements

The idea for this paper came whilst preparing a talk for a conference on “Biological invasions and evolutionary biology, stochastic and deterministic models” in Lyon, 2013. So I thank the organisers of the conference — Jean Berard, Vincent Calvez, and Gael Raoul — for both the conference and their invitation. I also thank Wayne Mallet and Jeremy Vanderwal for ongoing technical support around High Performance Computing. Funding for this work was provided by the Australian Research Council (DP1094646).

## 7 Biosketch

Ben Phillips is a QEII Research Fellow at the Department of Biosciences, University of Melbourne, Australia. His work focuses on contemporary evolution: how it plays out in space, and how it interacts with ecology. He has recently become fascinated by range edges and how evolution might affect whether, and how fast, range edges will shift under climate change. http://blphillipsresearch.wordpress.com

